# Genomic library of *Bordetella*

**DOI:** 10.1101/2022.01.20.475763

**Authors:** Sébastien Bridel, Valérie Bouchez, Bryan Brancotte, Sofia Hauck, Nathalie Armatys, Annie Landier, Estelle Mühle, Sophie Guillot, Julie Toubiana, Martin C.J. Maiden, Keith A. Jolley, Sylvain Brisse

## Abstract

**Background:** The re-emergence of whooping cough and geographic disparities in vaccine escape or antimicrobial resistance dynamics, underline the importance of a unified definition of *Bordetella pertussis* strains. Understanding of the evolutionary adaptations of *Bordetella* pathogens to humans and animals requires comparative studies with environmental *bordetellae*.

**Methods:** We have set-up a unified library of *Bordetella* genomes by merging previously existing Oxford and Pasteur databases, importing genomes from public repositories, and developing harmonized genotyping schemes. We developed a genus-wide cgMLST genotyping scheme and incorporated a previous *B. pertussis* cgMLST scheme. Specific schemes were developed to define antigenic, virulence and macrolide resistance profiles. Genomic sequencing of 83 French *B. bronchiseptica* isolates and of *B. tumulicola, B. muralis* and *B. tumbae* type strains was performed.

**Results:** The public library currently includes 2,581 *Bordetella* isolates and their provenance data, and 2,084 genomes. The “classical Bordetella” (*B. bronchiseptica, B. parapertussis* and *B. pertussis*), which form a single genomic species (*B. bronchiseptica* genomic species, BbGS), were overrepresented (n=2,382). The phylogenetic analysis of *Bordetella* genomes associated the three novel species *B. tumulicola, B. muralis* and *B. tumbae* in a clade with *B. petrii* and revealed 18 yet undescribed species. A sister lineage of the classical *bordetellae*, provisionally named *Bbs* lineage II, was uncovered and may represent a novel species (average nucleotide identity with BbGS strains: ∼95%). It comprised strain HT200 from India, two strains of ‘genogroup 6’ from the USA and six clinical isolates from France; this lineage lacked *ptxP* and its fim2 gene was divergent. Within *B. pertussis*, vaccine antigen sequence types marked important phylogenetic subdivisions, and macrolide resistance markers (23S_rRNA allele 13 and fhaB3) confirmed the current restriction of this phenotype in China with few exceptions.

**Conclusions:** The genomic platform provides an expandable resource for unified genotyping of *Bordetella* strains and will facilitate collective evolutionary and epidemiological understanding of the re-emergence of whooping cough and other *Bordetella* infections.

**Data summary:** ***Bordetella* genomes list and accession numbers: Supplementary Table S4**

***Bordetella* genus phylogeny dataset (92 isolates):** https://bigsdb.pasteur.fr/cgi-bin/bigsdb/bigsdb.pl?db=pubmlst_bordetella_isolates&page=query&project_list=23&submit=1

***B. bronchiseptica* phylogeny dataset (213 isolates):** https://bigsdb.pasteur.fr/cgi-bin/bigsdb/bigsdb.pl?db=pubmlst_bordetella_isolates&page=query&project_list=24&submit=1

***B. pertussis* phylogeny (124 isolates):** https://bigsdb.pasteur.fr/cgi-bin/bigsdb/bigsdb.pl?db=pubmlst_bordetella_isolates&page=query&project_list=25&submit=1

iTOL interactive trees: https://itol.embl.de/shared/1l7Fw0AvKOoCF

## Introduction

*Bordetellae* are beta-proteobacteria that can be found in the environment and can cause infections in animals and humans. Of the 16 currently described *Bordetella* species, the medically most important taxa *B. pertussis* (*Bp*) and *B. parapertussis (Bpp)*, together with *B. bronchiseptica* (*Bbs)*, are referred to as the ‘classical *bordetellae*’ and belong to a single genomic species [1], hereafter named the *B. bronchiseptica* genomic species (BbGS) for convenience. *B. pertussis*, and more rarely *B. parapertussis*, cause whooping cough, characterized by its typical paroxysmal cough, and kill an estimated ∼140,000 children annually [2–4]. *Bp* and *Bpp* have evolved from sublineages of *Bbs* [5]. Whereas *Bp* and *Bpp* are human restricted, *B. bronchiseptica* has a broader ecological distribution and causes respiratory infections in a wide range of mammalian hosts including humans. Based on MLST [1] and genomic sequencing [4,6] two distinct *Bbs* groups were named *Bbs* complexes I and IV [1], the last one being more strongly associated to humans.

Other *Bordetella* species were described and can affect humans, although a few were so far only described as environmental. *B. holmesii* can be collected either from blood of septicemic patients or from nasopharyngeal samples of patients with pertussis-like symptoms [7–9]. *B. hinzii* is frequently carried in birds [10,11] and *B. avium* and *B. pseudohinzii* are respectively responsible of respiratory disease in poultry or birds [12] and in mice or wild rats [13]. *B. bronchialis, B. sputigena* and *B. flabilis* were described from respiratory samples of patients with cystic fibrosis [14,15], and *B. trematum* and *B. ansorpii* were found in infected wounds of immunocompromised patients [4,16,17]. Some *Bordetella* species have been found in environmental samples, such as *B. petrii* or the three recently described species *B. muralis, B. tumulicola* and *B. tumbae* [18–20]. In addition to these 16 current *Bordetella* species with standing in the prokaryotic taxonomy, additional *Bordetella* genogroups (as also named *Bordetella* genomospecies in INSDC bioproject PRJNA385118) were identified from patients with cystic fibrosis and may represent additional separate *Bordetella* species [21].

The diversity of lifestyles and medical importance of *Bordetellae* raise important questions about the origins and evolution of pathogenicity in this group [4]. Horizontal gene transfer (HGT) is likely to occur between *Bordetella* species and strains, as already observed for the O-antigen locus in *B. bronchiseptica* [22]. For example, the environmental species *B. petrii* has genomic islands with atypical G+C content [18] that were associated with the metabolic versatility of this species. The use of a unified database of genomes from all *Bordetellae* species would facilitate genomic analyses underlying the phenotypic diversity within this important bacterial genus.

Besides the large-scale evolution of *Bordetellae*, the population dynamics within *Bp* are an important topic of epidemiological surveillance, in light of vaccine-escape evolution and the possible emergence and global dissemination of antimicrobial resistance [23–25]. Major evolutionary events in vaccine antigen-related genomic features have been described, including within the promoter region of the pertussis toxin gene cluster (*ptxP)* and in the fimbriae gene *fim3* [23]. In most countries using acellular vaccines, isolates characterized by *ptxP*3 and either *fim3*-1 or *fim3*-2 alleles are currently largely predominant, whereas isolates of the ancestral *ptxP*1 genotype still predominate elsewhere, for example in China [24,26]. In addition, in countries that use acellular vaccines, an increasing proportion of *Bp* isolates are deficient for the production of the immunodominant surface protein pertactin, raising questions on future vaccine effectiveness [27]. The nomenclature of genotypic markers, sublineages and strain subtypes needs unification to facilitate collective studies of the global epidemiology and population dynamics in *Bp*.

Harmonization of the genotype nomenclature of bacterial pathogens may be achieved by the gene-by-gene approach called MLST (for multilocus sequence typing), in which allele numbers are uniquely attributed to each locus sequence variant. Bioinformatics platforms that centralize allele definitions and allow curation and open access to genotype nomenclature are critical resources for unified pathogen subtype definitions [28]. Until now, there have been two dedicated *Bordetella* genome sequence databases: Oxford University’s PubMLST (unpublished) and Institut Pasteur’s BIGSdb [29] platforms. This duality has led to nomenclatural confusion and suboptimal services to the user’s community. With the rapid developments of genomic epidemiology and biodiversity surveys, a common and optimized resource for *Bordetella* evolutionary studies and sequence variants naming is needed. Here, we aimed to set-up such a unified genomic platform and expand its features to enable addressing genus-wide evolutionary genomics questions and global tracking of important genotypes within individual species, and particularly *B. pertussis*.

## Methods

### DNA preparation and genomic sequencing

Genomic sequencing was performed for 83 French *B. bronchiseptica* isolates mainly (84 out of 89) of human origin collected between 2007 and 2020, and for the type strains of *B. tumulicola, B. muralis* and *B. tumbae*.

Isolates were grown at 36°C for 72 hours on Bordet-Gengou agar (Becton Dickinson, Le Pont de Claix, France) supplemented with 15% defibrinated horse blood (BioMérieux, Marcy l’Étoile, France), and sub-cultured in the same medium for 24 hours. Bacteria were suspended in physiologic salt to reach OD_650_ of 1, and 400 μL of the suspension were pelleted. Pellets were re-suspended in 100 μL of PBS 1X, 100 μL of lysis buffer (Roche), and 40 μL of proteinase K; heated at 65°C for 10 minutes and at 95°C for 10 minutes; and used for DNA extraction. Whole genome sequencing was performed using a NextSeq 500 system (Illumina, USA) at the Mutualized Platform for Microbiology of Institut Pasteur. For *de novo* assembly, paired-ends reads were clipped and trimmed with AlienTrimmer [30], corrected with Musket [31], merged (if needed) with FLASH [32], and subjected to a digital normalization procedure with khmer [33]. For each sample, remaining processed reads were assembled and scaffolded with SPAdes [34].

### Development of pan-genus *Bordetella* cgMLST scheme

A genus-wide core genome MLST scheme [35], called *Bordetella* cgMLST v1.0, was established alongside a genus-wide locus annotation system (BORD loci), using the principles published for the genus *Neisseria* [36]. Six finished and annotated *Bordetella* genomes were uploaded into the BIGSdb Oxford database [37]: *Bordetella pertussis* Tohama I [38]; *Bordetella pertussis* CS [39]; *Bordetella bronchiseptica* RB50 [38]; *Bordetella parapertussis* 12822 [38]; *Bordetella petri* 12804 [18]; and, *Bordetella avium* 197N [40]. Using the GenomeComparator module of BIGSdb, the greatest number of common genes (1469) was found when *B. bronchiseptica* RB50 was used as the reference genome (even though *B. petri* 12804 had the greatest number of annotated loci, 5023). Genes described as ‘hypothetical’ or ‘putative’ were removed, resulting in 1415 loci. All loci entered into the database were given a BORD number of the format BORD000000, where the last six digits corresponded to the BB0000 numbers used in the annotation of *B. bronchiseptica* RB50 [38]. Other annotations and known aliases were included in the locus descriptions for comparison. As with other genus-wide locus schemes, this scheme can be expanded simply by adding additional loci, to give an expandable catalogue of the genus pangenome.

### Merging of the Oxford and Pasteur databases

Two BIGSdb databases were originally designed separately for distinct purposes: while Oxford’s PubMLST database offered MLST, *Bordetella* cgMLST v1.0 (see above) and virulence genes schemes, the Pasteur database was designed for the sole initial purpose of *Bordetella pertussis* genotyping, with a Bp–specific cgMLST scheme and a Bp-virulence genes schemes [29].

To merge the data available in the two databases, we proceeded as follows. As per BIGSdb dual design, isolates genomes and provenance data were imported into the “isolates” database, whereas allelic definitions of MLST, cgMLST and virulence genes were imported into the “seqdef” database.

Regarding the isolates database, we first dumped the Oxford database and uploaded it on the Institut Pasteur server. Second, we imported Pasteur isolates into this new database. To facilitate the understanding of historical origins of each entry, isolates identification numbers (BIGSdb ID number) were defined as follows: isolates from the original Oxford database were numbered from 1 to 1,914 as in the original database (*i*.*e*., their identification numbers were untouched). Second, the isolates from the original Pasteur database were numbered from 10,000 to 12,214 (original Pasteur identification number + 10000). We also completed the comment fields for these isolates, with the old identification number being added with the suffix word “Pasteur”. We added “putative duplicate” in this comment field if the corresponding isolate name was present in both original databases. We also added a “duplicate number” field in the isolates database. If two or more isolates were identified as the same strain, they were attributed the same duplicate number, consecutively. Duplicated data identification was based on a combined analysis of metadata and phylogeny tree reconstruction.

At time of merging and closure, the Oxford database comprised 1,914 isolates entries, 57 of which were private, whereas the Pasteur database comprised 2,180 entries, 2,009 of which were private. As of January 7^th^, 2022, the platform resulting from the merger comprises 2,581 public isolates entries, and 4,853 isolates in total when considering private entries.

### cgMLST schemes

Two core genome MLST schemes are available. The first scheme, *Bordetella* cgMLST v1.0 (hereafter referred to as *cgMLST_genus)*, was initially defined and hosted in the Oxford platform and was designed to be applicable for the entire *Bordetella* genus. The second scheme, called *cgMLST_pertussis*, was originally hosted in the Pasteur database and was built for *B. pertussis* isolates genotyping only [29]. Note that only the latter scheme has attached cgST definitions, *i*.*e*., unique cgST identifiers are attached to each distinct allelic profile. Both cgMLST schemes are available in the merged library.

### Genotyping schemes for vaccine antigens and virulence-associated genes

In both original databases, virulence-associated gene schemes had been defined separately. We choose to keep all loci from the virulence scheme designed in Oxford and to add some additional relevant loci. These loci were then grouped into different schemes comprising either Bp vaccine antigens (*fim2, fim3, ptxP, ptxA, ptxB, ptxC, ptxD, ptxE, fhaB*-2400_5550), T3SS genes (*bopB, bopD, bopN, bsp22, bteA*), autotransporters (*bapC, brkA, tcfA, prn, vag8*), other toxins (*cyaA, dnt*) or phase biology genes (*bipA, bvgA, bvgS*). We decided to include the *prn* locus in the autotransporter scheme rather than in the *Bp* vaccine antigens scheme, because prn-negative isolates would not allow the definition of sequence types (ST) for the vaccine antigens scheme.

To make analyses of virulence-associated and vaccine antigen genes more user-friendly, common gene names used in the literature were used as locus identifier, instead of the locus tags that were initially used in the Oxford database (*e*.*g*., BORD005020 was replaced by fim2 and BORD005021 by fim3). In addition, for five loci common to the original Pasteur and Oxford databases (*i*.*e. fim2, fim3, ptxA, ptxP* and *tcfA*), the allele numbering was re-defined so that the alleles of each locus would match the nomenclature found in the literature for *B. pertussis* alleles; the correspondence between the original allele numbering system and the new one is defined in **Table S1**. Subsequently, the numbering of alleles was simply incremented as new genomes scans led to novel allele definitions.

### Macrolide resistance

The 23S rRNA allele table was incremented with a sequence defined as allele 13, which carries the only mutation described so far as being associated with macrolide resistance [41]. To facilitate the detection of putative macrolide resistant isolates and their lineage [41,42], we built a scheme named “macrolide resistance” that includes the following loci: 23S_rRNA, fhaB (full) and four *fhaB* fragments. As the *fhaB* locus, which has a full-length size of 10,773 bp, is often fragmented in the genomic assemblies, three smaller fragments were designed to cover evenly the *fhaB* sequence: fhaBx-1_3193, fhaBy-3190_7183, fhaBz-7180_10773; the ranges in these locus names correspond to the begin and end positions of the three fragments. A fourth fragment was designed to cover the region comprising 2 SNPs present individually in *fhaB2* and *fhaB3* [24] : fhaB-2400_5550. The allele numbering for *fhaB* fragments was defined so that the alleles of each locus would match the nomenclature found in the literature, ie. alleles *fhaB1* and *fhaB2* correspond to those defined by van Loo [43], and *fhaB3* to the mutation C53301T [24]. Allele *fhaB3* of full size locus and restricted locus (fhaBy-3190_7183 and fhaB-2400_5550 )is associated with *ptxP1* isolates originating from China which exhibit a macrolide resistance phenotype [24].

### Genome scanning for defined loci

Reference genomes were selected from either type-strain, reference genomes from literature and/or RefSeq genomes or most complete genomes available for each species (*e*.*g*., type-strain 18323 and reference strain Tomaha for *B. pertussis*). All these genomes are grouped into the public project “Bordetella Genus Phylogeny”. We selected two representatives for each species, when available. All these genomes are found in the “Bordetella phylogeny” public project in the BIGSdb genome library. For the *cgMLST_genus* scheme, missing alleles were captured by relaxing the scanning parameters to 70% identity and 70% alignment in order to capture alleles from all *Bordetella* species, using the reference strains. New captured alleles were defining as type alleles. Then, all future detection of new alleles were based on 90% identity, 90% alignment parameters using defined type alleles as queries. Criteria for loci associated with virulence were stricter: only new alleles sharing 90% identity, 90% alignment with a type allele and with a complete CDS were recorded (excluding the *ptxP* promoter locus and the *fhaB* fragment fhaB-2400_5500). These strict parameters were used to minimize the detection of paralogous sequences and to exclude decaying alleles.

For each species, we defined one or two reference strains taken either from literature or from the NCBI RefSeq database (**Table S2)**. Type alleles were defined from each reference isolate in order to constrain the search space so as to ensure future consistency of allele definitions. The definition of new alleles was subject to constraining parameters. To detect alleles, BLASTN thresholds were set to 90% of identity, 90% of alignment length coverage, and considering only the complete coding sequences (except for the *ptxP* locus promoter and the 23S rRNA gene, which are not protein CDSs). These strict parameters were used to minimize the detection of paralogous sequences.

### Phylogenetic analyses

Phylogenetic analyses were performed based on concatenated alignments of individual gene loci defined in the database. We used an in-house script which combines IQtree (IQ-TREE 2.0.6, model substitution GTR+G+I) and Gubbins (2.4.1, default parameters) [44,45]. Phylogenetic trees were drawn and annotated using iTOL and FigTree [46,47].

To include the breadth of currently sampled *Bordetella* diversity, we first downloaded all genomic sequences from public databases and ran a rapid distance based phylogenetic analysis. From this, we retained all representative genomes of unique branches and down-sampled shallow branches that were represented multiple times. The resulting dataset included the putative novel species initially described as genogroups by Spilker *et al*. on the basis of *nrdA* gene sequences [21]. The *Bordetella* diversity dataset is publicly available from the genomic library as project *Bordetella* genus phylogeny (https://bigsdb.pasteur.fr/cgi-bin/bigsdb/bigsdb.pl?db=pubmlst_bordetella_isolates&page=query&project_list=23&submit=1).

To analyze the phylogenetic structure within the BbGS, we selected 204 isolates: 177 *B. bronchiseptica* representative of the clonal complexes I and IV [1] and 9 others that did not belong to these complexes. This dataset was completed with strains from *B. pertussis* (n=9) and *B. parapertussis* (n=9), selected to represent the known diversity within these two taxa: for Bp, lineage I (strains B0887 and 18323) and lineage II (which includes *ptxP1* strains as including Tohama, and *ptxP3* strains) [23]; and for *B. parapertussis*, the ovine lineage (strain Bpp5) and the human lineage reference strain 12822. A public project encompassing all selected genomes is available in the BIGSdb genomic library platform (project name: *B. bronchiseptica* phylogeny; URL: https://bigsdb.pasteur.fr/cgi-bin/bigsdb/bigsdb.pl?db=pubmlst_bordetella_isolates&page=query&project_list=24&submit=1).

In the same way, 124 *Bp* isolates selected to represent *Bp* genomic diversity were used to analyze the phylogenetic position of macrolide-resistant isolates. To this aim, we selected isolates representative of main *ptxP* branches (*ptxP* alleles 1, 2, 3, 19 and 21) and all genomes of isolates known to be resistant to erythromycin (n=51). The dataset is publicly available from the genomic library as project *B. pertussis* phylogeny (https://bigsdb.pasteur.fr/cgi-bin/bigsdb/bigsdb.pl?db=pubmlst_bordetella_isolates&page=query&project_list=25&submit=1).

## Results

### Genomic library contents

After merging Oxford and Pasteur databases, importing publicly available genomes, and adding three genomes of recently described species *B. muralis* (CIP111920^T^), *B. tumulicola* (CIP111922 ^T^) and *B. tumbae* (CIP111921 ^T^) that we sequenced, the genomic library comprised representatives all 16 *Bordetella* species currently described in the Prokaryotic taxonomy. The number of genomes varied across species, the most represented ones being the classical *Bordetella* species *B. pertussis* (n = 1,567), *B. bronchiseptica* (n = 618) *and B. parapertussis* (n = 197). *Bordetella holmesii* (n = 59) and the other species were represented by fewer genomes, including some by only one genome, such as *B. flabilis* or *B. sputigena*.

### Phylogenetic analysis of the genus

A phylogenetic analysis of the 16 currently described *Bordetella* species and putative novel *Bordetella* species was performed, based on the alignments of the sequences of the genus-wide cgMLST scheme loci (**Figure 1)**. As expected, the three classical *Bordetella* species were grouped in a common clade. A second clade comprised *B. holmesii, B. avium, B. hinzii* and *B. trematum*, consistent with Linz et al. [4], and also *B. pseudohinzii* and *Bordetella* genogroups 12, 5 and 1 [21]. A third major clade comprised *B. petrii* and *Bordetella* genogroups 7, 2 and 4. The three novel species *B. muralis, B. tumbae* and *B. tumulicola* were part of a single branch distantly associated to the *B. petrii* clade, consistent with 16S based phylogenetic analyses [20]. *B. flabilis, B. sputigena* and *B. bronchialis* were found in a fourth, more distant group together with genogroups 8, 9, 10 and 11. Finally, the earliest branching *Bordetella* species was *B. ansorpii*, as previously observed [4], and it was associated with genogroup 13.

**Figure 1:**
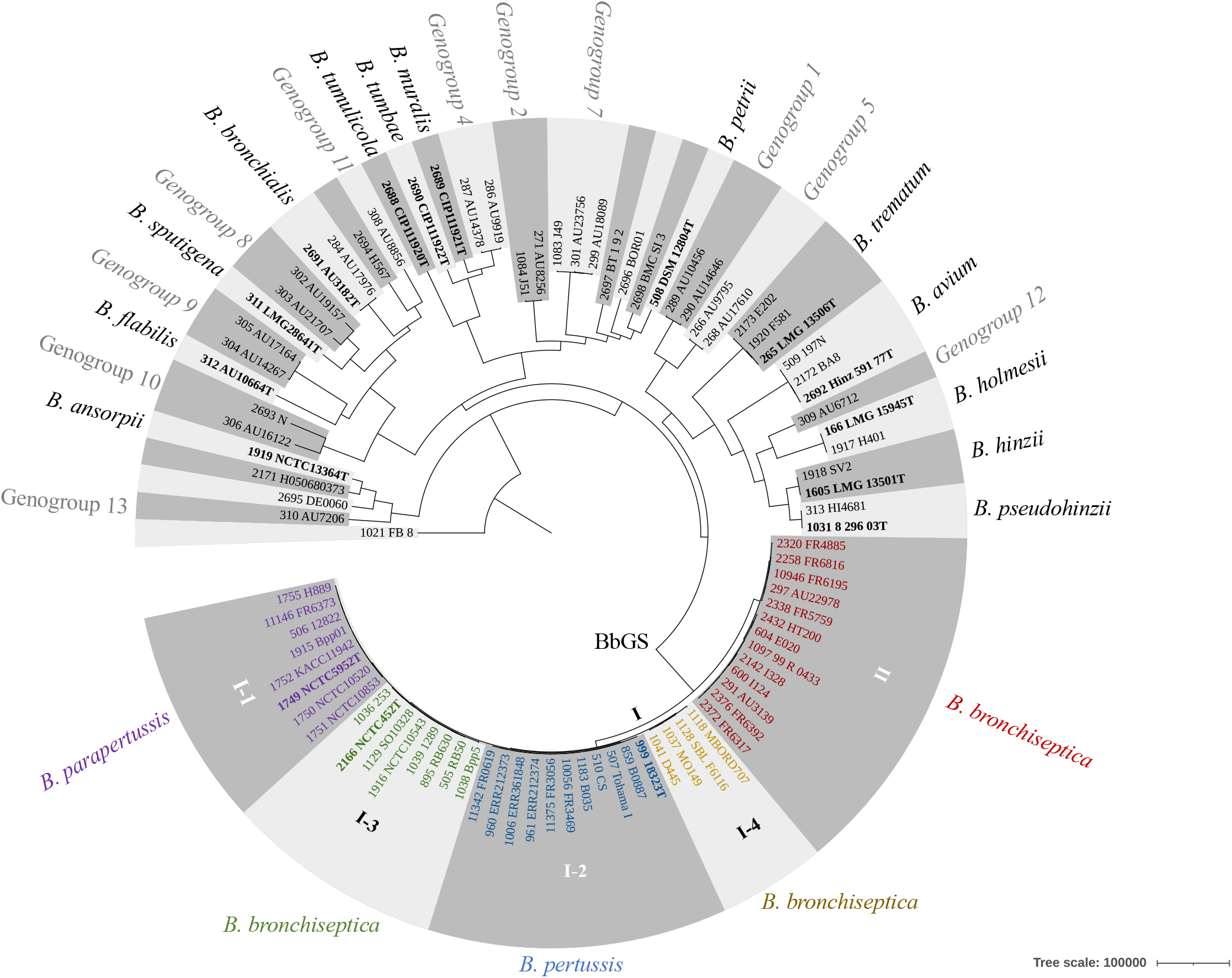
Phylogenetic analysis of the *Bordetella* genus. The phylogenetic tree was obtained based on the concatenated multiple sequence alignments of the 1,415 core gene sequences from the *cgMLST_genus* scheme; recombination was accounted for using Gubbins. The tree was rooted on the branch leading to strain FB-8, which was the earliest branching genome in a phylogenetic analysis that included the external group *Ralstonia solanacearum* strain IBSBF1503 (Figure S4). Leaves are labelled with the identifier of the strain in the BIGSdb database, followed by the strain name.

The ANI metric is widely used as a guide to define novel bacterial species [49]. ANI values between representative genomes of the *Bordetella* genus (**Table S2; Figure S1**) showed that most currently described species differ by >5% nucleotide positions, with the well-known exception of BbGS members, which present ANI values between 98.3 and 99.4%. Here, we note that a large number of some previously attributed species names would need reevaluation; for example, *B. petrii* type strain and ‘*B. petrii’* strain J51 had only 85.14% ANI, indicating that the latter should be considered as belonging to a distinct species, which remains to be defined. Likewise, ‘*B. ansorpii*’ strain H050680373 had an ANI of 93.48% with *B. ansorpii* type strain NCTC13364^T^ **(=** SMC-8986^T^). Some species correspond to previously defined genogroups, as observed for *B. bronchialis* and genogroup 3 (isolates AU3182^T^ and AU17976), for *B. sputigena* and genogroup 14 (isolates LMG28641^T^, AU21707 and AU19157) and for *B. flabilis* and genogroup 15 (isolate AU10664^T^).

### Phylogenetic diversity within *Bordetella bronchiseptica*

A phylogenetic tree of *Bbs* isolates, including 83 isolates from France sequenced for this study, was inferred from the *cgMLST_genus* gene sequences (**Figure 2 & Figure S2**) and revealed a primary separation of *Bbs* diversity into two deep lineages. We here define the minor one as lineage II of *B. bronchiseptica*. Lineage II comprised nine isolates, one of which is HT200 from a natural spring water in India and was previously recognized as being atypically distant from other *Bbs* isolates [50]. *Bbs* lineage II isolates from the USA (AU22978 et AU3139) were previously described as genogroup 6 and had been collected from patients with cystic fibrosis [21]. The remaining lineage II isolates were six human clinical isolates from France, collected from adults (mean age = 71.7 years) displaying pulmonary infections. These observations clearly establish the pathogenic potential of *Bbs* lineage II. In turn, *Bbs* lineage I comprised four sublineages, which we define as sublineages I-1 to I-4 in order to mirror the previous 7-gene MLST-defined clonal complexes I (here, sublineage I-1), II (Bp, sublineage I-2), III (Bpp, sublineage I-3) and IV (sublineage I-4). The ANI values between *Bbs* lineage II and the four sublineages of lineage I ranged from 95.09% to 95.72%, indicating that *Bbs* lineage II may be considered as a species distinct from lineage I, which itself equates to the BbGS or ‘classical *bordetellae*’.

**Figure 2:**
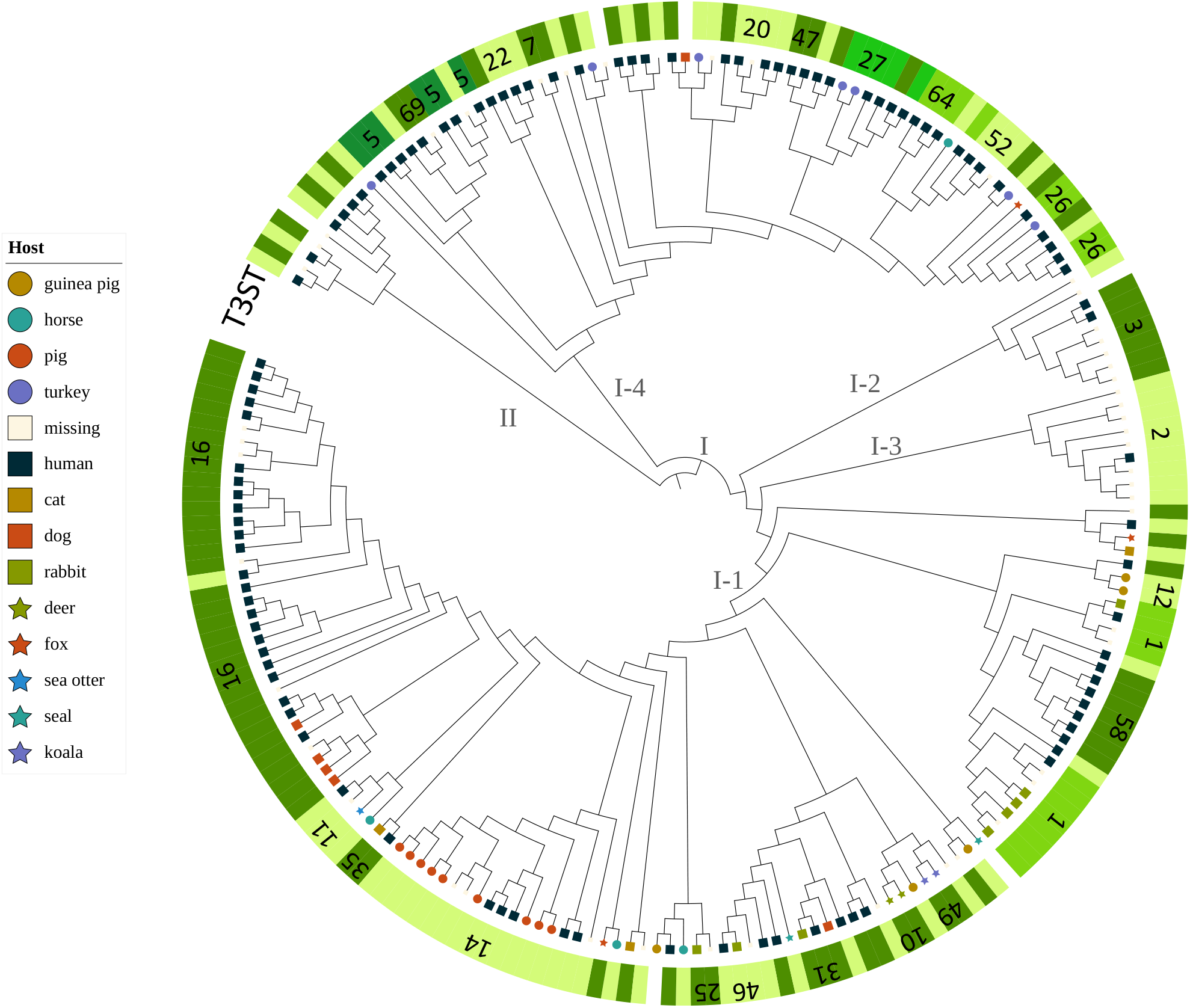
Cladogram of *Bordetella bronchiseptica*. The analysis was performed with 186 *B. bronchiseptica* genomes, and representatives of the phylogenetic diversity of *B. pertussis* (9 genomes) and *B. parapertussis* (9 genomes). The recombination-purged concatenated multiple sequence alignment of 1,415 core gene loci (*cgMLST_genus* scheme) was used. The tree is rooted on lineage II, which is the most divergent clade. Branch lengths were not used in order to ease readability of groups and isolates; see Figure S5 for the corresponding phylogram. For each isolate, the host is represented using a leaf symbol, where available (see key in Figure S5). The numbers corresponding to T3SS sequence types are indicated along the external circle around the tree; only the identifiers of main T3STs are indicated.

The independent origins of *Bpp*_hu_ and *Bpp*_ov_ have been debated [1,6]. We note that Bpp5, the reference strain of ovine *B. parapertu*ssis, was associated with the earliest branching *Bbs* sublineage I-1, rendering *B. parapertussis* paraphyletic (**Figures 2 & S2**). A separate phylogenetic analysis performed using the *cgMLST_pertussis* scheme, instead of the genus-wide scheme, provided a nearly identical phylogenetic tree of *Bbs* isolates (data not shown). However, we recommend using the *cgMLST_genus* scheme for *Bbs* phylogenetic analyses, as a lower number of uncalled alleles was observed than for the *cgMLST_pertussis* scheme (**Figure S3**).

### Genotyping of vaccine antigen and virulence-associated genes of *Bordetella bronchiseptica*

The allele diversity and presence/absence of vaccine antigens or virulence-associated loci were investigated within *Bbs*. For convenience, the loci were grouped into schemes labeled as Bp-vaccine antigens, Autotransporters, Phase genes, Other Toxins and T3SS.

T3SS genes are known to be present within the BbGS [6]. However, isolates of the classical *Bordetella* species differ in the levels of expression of their T3SS *in vitro* [51]: although *Bbs* and ovine *Bpp* are able to produce T3SS effectors such as BteA, in human adapted *Bp* and *Bpp*, T3SS expression is blocked at post-transcriptional level [3,51]. The insertion of an alanine in position 503 of BteA from *B. pertussis* leads to the attenuation of *B. pertussis* cytotoxicity [52]. This insertion was found in 8 of 11 strains of *Bbs* sublineage II-2 (*Bp*), corresponding to alleles 1 and 6. We devised a nomenclature of sequence types based on the combination of alleles at the loci *bteA, bopB, bopD, bopN* and *bsp22*; these are referred to as T3STs. For T3SS genes *bteA, bopB, bopD, bopN* and *bsp22*, most alleles were specific of (sub)lineages with some minor exceptions. T3STs were largely specific for *Bbs* sublineages (**Figure 2 & S5)**. Sublineage I-1 comprised 30 T3STs, the main ones being type 16 and type 14, and sublineage I-4 comprised 33 T3ST types. Hypervirulent *Bbs* strains have been defined previously based on an enhanced activity of the T3SS, for example strain 1289 (BIGSdb ID: 1039; T3ST: 19) in sublineage I-1 or strain Bbr77 (BIGSdb ID: 1040; T3ST: 20) in sublineage I-4 [53,54]; however, they do not share the same T3ST. Interestingly, sublineage I-2 (*Bp*) isolates all belonged to T3ST-3, whereas isolates of sublineage I-3 (*Bpp*) all had T3ST-2, except for strain Bpp5, the ovine representative, with T3ST-18. T3ST were defined for 8 of 9 *Bbs* isolates of lineage II, all being distinct. The T3ST nomenclature therefore provides a complementary classification system of *B. bronchiseptica*, useful for distinction of subtypes within the two major sublineages of lineage I, and within lineage II.

For most other loci, alleles were specific for either *Bbs* lineage I or II, or for unique sublineages within lineage I (see details in the **Supplementary text** and in **Figure S5**). Consistent with its large (∼5%) genomic divergence from the BbGS, alleles observed within *B. bronchiseptica* lineage II were unique for this lineage. In addition, even though only few isolates have been sampled so far for this lineage, we noted a high diversity of alleles indicating that it forms a genetically diverse group, suggesting phenotypic variability. Some loci including *fim2, ptxP, dnt, tcfA, vag8, prn* or *bipA* had no allele tagged according to our stringent criteria. When relaxing these criteria, we noted that *ptxP, prn and dnt* were absent in *Bbs* lineage II, whereas *fim2, bipA* and *tcfA* presented partial matches (< 90%) with existing alleles in almost all isolates.

### Allele diversity and macrolide resistance in *B. pertussis*

A phylogeny of *Bp* isolates, which were selected with a focus on macrolide resistance, was built using the 2,038 loci of *cgMLST_pertussis* scheme (**Figure 3)**. Three isolates of the early *ptxP2* branch correspond to *Bp* lineage II-a as defined by Bart *et al*. [23]. The remaining isolates fell in the *ptxP1* and *ptxP3* branches, which represent the main subdivisions of *Bp* lineage IIb.

**Figure 3:**
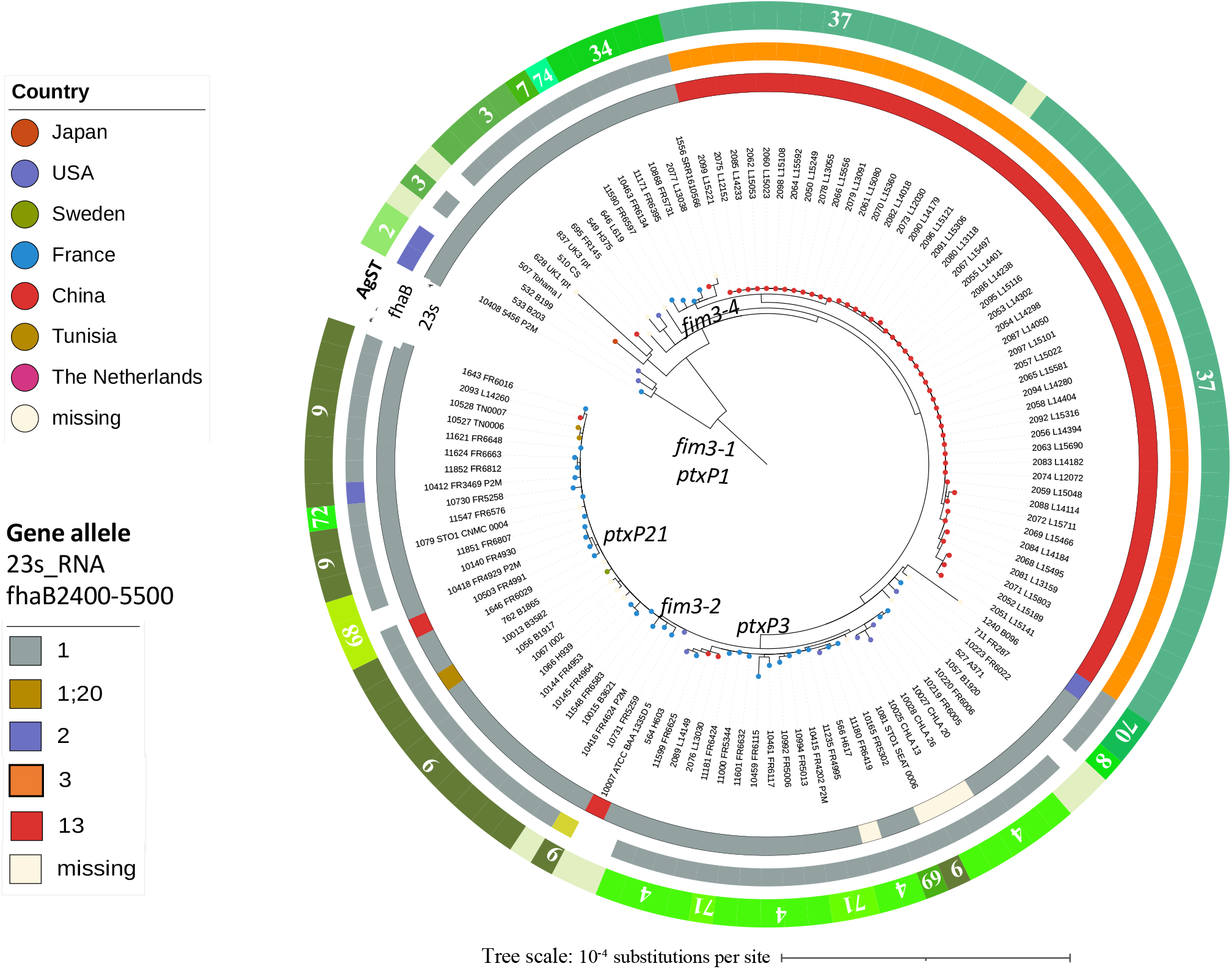
Phylogenetic analysis of *Bordetella pertussis*. The phylogenetic tree was obtained based on the recombination-purged concatenated multiple sequence alignments of the 2,038 core genome loci of the *cgMLST_pertussis* scheme. The distribution of macrolide resistance is indicated. The three outer circles represent (from innermost): *23S_rRNA* alleles, *fhaB* alleles and the vaccine antigen sequence types (AgST).

*Bp* isolates were screened with the *macrolide resistance* scheme to detect alleles associated with this phenotype. Allele 13 of the 23S_rRNA locus is specific for erythromycin resistant *B. pertussis* isolates, whereas alleles 1, 2 and 20 correspond to susceptible ones. Allele 13 was found in 46 of 51 isolates from China [24] and in two isolates previously described as erythromycin resistant: one from USA (ATCC:BAA-1335D-5, [42]) and one from France (FR4991, [41]). When checking our entire library, which contained 1,567 *Bp* isolates, no additional resistant isolate was found. In a previous study, Xu *et al*. [24] showed that allele *fhaB*3 characterized by the mutation C5330T was found in a unique branch of the phylogenetic tree of *Bp*, and that this branch corresponds to macrolide-resistant isolates from China. Here, macrolide-resistant isolates from China shared the same allele for *fhaB* (full): fhaB-39. Only one exception was apparent, for isolate L14404 which had a 15-nucleotide insertion at position 6959, defined as allele *fhaB*-40. In our library, the alleles observed in isolates of the *fhaB*3 branch were defined as fhaBy-3190_7183 allele 3, fhaB-2400_5550 allele 3 and 23S rRNA allele 13 (following the nomenclature from literature [24]). Whereas all isolates from China with 23S rRNA allele 13 have the fhaB3 allele, the opposite was not true, as some isolates (PT2019, ERR030030, ERR361878, SRR1610553) have the fhaB-2400_5550 allele 3, but 23S rRNA allele 1. Hence, the *fhaB3* allele marks a broader lineage, of which a subset of isolates possesses the 23S rRNA allele 13. As observed in the phylogenetic tree, macrolide-resistant *Bp* isolates from China all fell in a single branch (characterized by both fhaB2400_5500 allele 3 and 23S rRNA allele 13). The two other resistant isolates, collected in the USA and France, belonged to the *ptx*P3/*fim*3-2 branch and were not grouped together, indicating three independent evolutionary origins of macrolide-resistance in these isolates and the Chinese ones. The macrolide resistant isolates from the USA and France both displayed allele 1 for fhaB2400_5500 locus and allele 13 for 23s_rRNA.

We also defined a *Bp* vaccine antigens scheme, which groups the loci *ptxP, ptxABCDE, fim2, fim3* and *fhaB2400_5550*. Bp vaccine antigens sequence types (BPagST) were defined for this scheme, corresponding to the unique combinations of alleles at the individual loci. These STs were distributed along the *Bp* tree (**Figure 3)**. For example, BPagST34 marked the sublineage with allele *fim3*-4, whereas BPagST9 corresponded to most isolates of the *fim3*-2 branch. BPagST37 (allele 3 for *fhaB2400_5550* locus) corresponded to macrolide resistant *fhaB3* lineage of Xu *et al*. [24]. The macrolide-resistant isolate from France was characterized by BPagST68, whereas no BPagST could be determined for the macrolide resistant isolate from the USA, because its allele at the *fim3* locus was not defined due to a frameshift. We suggest that the BPagST sequence types will be useful for future tracking of specific sublineages of *Bp*.

Loci variation within other schemes (Other toxins, Autotransporters, T3SS and Phase biology schemes) are detailed in the **Supplementary Text**.

## Discussion

We have set-up a unified genomic platform, available at https://bigsdb.pasteur.fr/bordetella/, by merging the data from the two previously existing PubMLST and Pasteur BIGSdb-*Bordetella* databases. This merger has enabled allelic nomenclature unification and has led to reassemble on the same platform, genotyping schemes that will facilitate the study of *Bordetella* diversity at different phylogenetic breadths. We have also expanded the genomic library with public sequences and sequenced genomes from the French NRC. The resulting unified platform for *Bordetella* genotyping will ease the future curation and expansion of *Bordetella* genomic data by a community of users and curators.

Two complementary cgMLST schemes are available on the genomic platform. While the *cgMLST_genus* scheme should be preferred for genus-wide phylogenetic analyses and within individual species including *B. bronchiseptica*, the *cgMLST_pertussis* scheme comprises more loci that are conserved within *Bp*, and its added discriminatory power will be useful for strain identification and comparison within *Bp*. Recently, a whole-genome MLST (wgMLST) scheme comprising even more loci (n =3,506; 1,822 of which are common with our *cgMLST_pertussis* scheme) was proposed for Bp subtyping, and concordant results were obtained for classification of outbreak and sporadic isolates from the USA using the cgMLST and wgMLST schemes [55]. As the later provides slightly higher discrimination among Bp isolates than the *cgMLST_pertussis* scheme, it might provide valuable additional genotyping information for local transmission investigations, and may be incorporated into the unified platform in the future.

We provided examples of applications of the genomic library and its associated genotyping schemes, from the analysis of the genus-level diversity of species, to strain genotyping within *Bp*. Regarding the phylogenetic structure of *Bordetella*, we provide a complete picture of the relationships among the 16 *Bordetella* species that have current standing in the Prokaryotic taxonomy, and confirm the existence of approximately the same number of additional putative species ([21]; PRJNA385118). This work shows that the taxonomy of the *Bordetella* genus will need future updates, and the unified *Bordetella* genomic library should facilitate the recognition of strains belonging to the same putative novel *Bordetella* taxa, thereby enhancing knowledge of their microbiological, ecological or pathogenic properties.

An important finding of this work is the identification of a novel *B. bronchiseptica* lineage, which we provisionally named *Bbs* lineage II. This lineage is clearly distinct from the classical *Bordetella*, which we therefore propose to define strictly as the BbGS, *i*.*e*., encompassing the previously described *Bbs* lineages I-1 and I-4 together with *Bp* and *Bpp*. The average ANI between *Bbs* lineage II and the BbGS members is borderline regarding the species definition threshold (95-96%), and the relevance of elevating *Bbs* lineage II as a taxonomic species or subspecies separate from *B. bronchiseptica* remains to be defined. All but one isolate from this lineage were collected from humans, demonstrating the pathogenic potential of *Bbs* lineage II, as for lineages I-1 and I-4. Virulence factors content and vaccine antigens sequence variants are unique to lineage II. Whether lineage II has distinctive epidemiology or pathogenic features should be the subject of future studies.

A rarely isolated subgroup of *Bpp* causes pneumonia in sheep and has been referred to as *Bpp*_ov_ in order to mark its distinctness from *Bpp*_hu_, the human *Bpp* [56]. *Bpp*_ov_ remains an under-studied lineage within the BbGS. So far, very few isolates of ovine *Bpp* have been reported, mainly from New Zealand and Scotland, and all were collected before 2006 [56–58], suggesting that this lineage is very rare, if at all still circulating. Bpp5 is the only *Bpp*_ov_ representative for which a genome sequence is currently available. More genomes of *Bpp*_ov_ would be needed to define the phylogenetic position and unique genomic features of this group within the BbGS.

*Bp* is the medically most important *Bordetella* species, being the most frequent cause of important infections in humans. The unified *Bordetella* genomic platform provides specific tools for characterizing genes linked to *Bp* vaccine antigens and virulence. These were organized into several genotyping schemes that might be used separately for specific biological or epidemiological questions. Where available, we defined allele identifiers in accordance with their literature numbering. This will disambiguate, in particular, the genotyping of vaccine antigens within *Bp*. The population of *Bp* has been noted to evolve in response to the selective pressure exerted by vaccination, in particular in countries using the acellular vaccines [23,59]. The proposed vaccine schemes will help keeping track of the circulating strains and defining the prevalence of particular sublineages, including those that are deficient for pertactin production [27,60,61].

We also created a genotyping scheme dedicated to macrolide resistance, the possible dissemination of which is an important point of public health concern. To enable easy identification of the macrolide-resistant sublineage that is currently highly prevalent in China, we defined this scheme as combining the causative allele 23S rRNA with the associated marker *fhaB*3. This scheme thus provides a convenient way to distinguish ‘out of China’ dissemination of the fhaB3 sublineage, from parallel evolution of macrolide resistance in other locations.

An important current limitation of the genomic library is its contents imbalance towards genomic sequences from acellular vaccine-using high-income countries, as observed for other pathogens [62]: our database contained only few genomes from Africa (0.2 %), Asia (1.9 %) or Oceania (2.1 %) as compared to those from North America (13.6 %) or Europe (58.7 %). In the future, efforts should be made to generate and incorporate more genomic sequences from world regions that are currently underrepresented.

In conclusion, the unified *Bordetella* genomic library was built with the intention to facilitate and harmonize the genotyping of strains of this important bacterial genus, and should represent a useful tool even for genomic epidemiology consortia unfamiliar with genomic analyses. It provides a uniformization of genetic variants designations, which will clarify the communication on genotypes and will enable a collective understanding on the biodiversity and epidemiology of *Bordetella*. We provide a timely solution for genomic studies of *Bordetella* pathogens, and in particular *Bp*, the reemergence of which is partly caused by evolutionary and epidemiological dynamics of public health concern, including vaccine escape and antibiotic resistance emergence.

## Author contributions

**Conceptualisation, Supervision**: SBrisse designed and coordinated the study. **Methodology, software, validation and data curatio**n: SBridel performed the merging the two databases into BIGSdb-Pasteur and the phylogenetic analyses, as well as extensive databases curation (metadata and duplicated data cleaning). VB performed in-depth analyses of virulence and antigen loci sequences and established the literature-based allelic nomenclature. SH, KAJ and MCJM created the *Bordetella* cgMLST v1.0 scheme. KAJ & MCJM provided the data from BIGSdb Oxford and developments of the BIGSdb platform. NA, AL, SG, JT and EM analyzed *Bordetella* strains at the French NRC and performed genomic sequencing. BB provided support for the BIGSdb-Pasteur platform deployment and maintenance. **Writing-Original draft Preparation, Review and Editing**: SBridel, VB and SBrisse wrote the original draft. All authors contributed to and approved the final version of the manuscript.

## Acknowledgments

We thank Julien Guglielmini (Institut Pasteur) for help with the BIGSdb-Pasteur database configuration.

## References

[1] Diavatopoulos DA, Cummings CA, Schouls LM, Brinig MM, Relman DA, Mooi FR. Bordetella pertussis, the Causative Agent of Whooping Cough, Evolved from a Distinct, Human-Associated Lineage of B. bronchiseptica. PLOS Pathog 2005;1:e45. https://doi.org/10.1371/journal.ppat.0010045.

[2] Heininger U, Stehr K, Schmitt-Grohé S, Lorenz C, Rost R, Christenson PD, et al. Clinical characteristics of illness caused by Bordetella parapertussis compared with illness caused by Bordetella pertussis. Pediatr Infect Dis J 1994;13:306–9. https://doi.org/10.1097/00006454-199404000-00011.

[3] Mattoo S, Cherry JD. Molecular Pathogenesis, Epidemiology, and Clinical Manifestations of Respiratory Infections Due to Bordetella pertussis and Other Bordetella Subspecies. Clin Microbiol Rev 2005;18:326–82. https://doi.org/10.1128/CMR.18.2.326-382.2005.

[4] Linz B, Ma L, Rivera I, Harvill ET. Genotypic and Phenotypic Adaptation of Pathogens: Lesson from the Genus Bordetella. Curr Opin Infect Dis 2019;32:223–30. https://doi.org/10.1097/QCO.0000000000000549.

[5] Diavatopoulos DA, Cummings CA, Schouls LM, Brinig MM, Relman DA, Mooi FR. Bordetella pertussis, the causative agent of whooping cough, evolved from a distinct, human-associated lineage of B. bronchiseptica. PLoS Pathog 2005;1:e45. https://doi.org/10.1371/journal.ppat.0010045.

[6] Park J, Zhang Y, Buboltz AM, Zhang X, Schuster SC, Ahuja U, et al. Comparative genomics of the classical Bordetella subspecies: the evolution and exchange of virulence-associated diversity amongst closely related pathogens. BMC Genomics 2012;13:545. https://doi.org/10.1186/1471-2164-13-545.

[7] Weyant RS, Hollis DG, Weaver RE, Amin MF, Steigerwalt AG, O’Connor SP, et al. Bordetella holmesii sp. nov., a new gram-negative species associated with septicemia. J Clin Microbiol 1995;33:1–7. https://doi.org/10.1128/jcm.33.1.1-7.1995.

[8] Pittet LF, Emonet S, Schrenzel J, Siegrist C-A, Posfay-Barbe KM. Bordetella holmesii: an under-recognised Bordetella species. Lancet Infect Dis 2014;14:510–9. https://doi.org/10.1016/S1473-3099(14)70021-0.

[9] Harvill ET, Goodfield LL, Ivanov Y, Smallridge WE, Meyer JA, Cassiday PK, et al. Genome Sequences of Nine Bordetella holmesii Strains Isolated in the United States. Genome Announc 2014;2:e00438–14. https://doi.org/10.1128/genomeA.00438-14.

[10] Funke G, Hess T, von Graevenitz A, Vandamme P. Characteristics of Bordetella hinzii strains isolated from a cystic fibrosis patient over a 3-year period. J Clin Microbiol 1996;34:966–9. https://doi.org/10.1128/jcm.34.4.966-969.1996.

[11] Collercandy N, Petillon C, Abid M, Descours C, Carvalho-Schneider C, Mereghetti L, et al. Bordetella hinzii: an Unusual Pathogen in Human Urinary Tract Infection. J Clin Microbiol 2021;59:e02748–20. https://doi.org/10.1128/JCM.02748-20.

[12] Raffel TR, Register KB, Marks SA, Temple L. Prevalence of Bordetella avium infection in selected wild and domesticated birds in the eastern USA. J Wildl Dis 2002;38:40–6. https://doi.org/10.7589/0090-3558-38.1.40.

[13] Ivanov YV, Linz B, Register KB, Newman JD, Taylor DL, Boschert KR, et al. Identification and taxonomic characterization of Bordetella pseudohinzii sp. nov. isolated from laboratory-raised mice. Int J Syst Evol Microbiol 2016;66:5452–9. https://doi.org/10.1099/ijsem.0.001540.

[14] Vandamme PA, Peeters C, Cnockaert M, Inganäs E, Falsen E, Moore ERB, et al. Bordetella bronchialis sp. nov., Bordetella flabilis sp. nov. and Bordetella sputigena sp. nov., isolated from human respiratory specimens, and reclassification of Achromobacter sediminum Zhang et al. 2014 as Verticia sediminum gen. nov., comb. nov. Int J Syst Evol Microbiol 2015;65:3674–82. https://doi.org/10.1099/ijsem.0.000473.

[15] Spilker T, Darrah R, LiPuma JJ. Complete Genome Sequences of Bordetella flabilis, Bordetella bronchialis, and “Bordetella pseudohinzii.” Genome Announc 2016;4:e01132–16. https://doi.org/10.1128/genomeA.01132-16.

[16] Castro TR, Martins RCR, Dal Forno NLF, Santana L, Rossi F, Schwarzbold AV, et al. Bordetella trematum infection: case report and review of previous cases. BMC Infect Dis 2019;19:485. https://doi.org/10.1186/s12879-019-4046-8.

[17] Ko KS, Peck KR, Oh WS, Lee NY, Lee JH, Song J-H. New species of Bordetella, Bordetella ansorpii sp. nov., isolated from the purulent exudate of an epidermal cyst. J Clin Microbiol 2005;43:2516–9. https://doi.org/10.1128/JCM.43.5.2516-2519.2005.

[18] Gross R, Guzman CA, Sebaihia M, dos Santos VAPM, Pieper DH, Koebnik R, et al. The missing link: Bordetella petrii is endowed with both the metabolic versatility of environmental bacteria and virulence traits of pathogenic Bordetellae. BMC Genomics 2008;9:449. https://doi.org/10.1186/1471-2164-9-449.

[19] Le Coustumier A, Njamkepo E, Cattoir V, Guillot S, Guiso N. Bordetella petrii infection with long-lasting persistence in human. Emerg Infect Dis 2011;17:612–8. https://doi.org/10.3201/eid1704.101480.

[20] Tazato N, Handa Y, Nishijima M, Kigawa R, Sano C, Sugiyama J. Novel environmental species isolated from the plaster wall surface of mural paintings in the Takamatsuzuka tumulus: Bordetella muralis sp. nov., Bordetella tumulicola sp. nov. and Bordetella tumbae sp. nov. Int J Syst Evol Microbiol 2015;65:4830–8. https://doi.org/10.1099/ijsem.0.000655.

[21] Spilker T, Leber AL, Marcon MJ, Newton DW, Darrah R, Vandamme P, et al. A simplified sequence-based identification scheme for Bordetella reveals several putative novel species. J Clin Microbiol 2014;52:674–7. https://doi.org/10.1128/JCM.02572-13.

[22] Buboltz AM, Nicholson TL, Karanikas AT, Preston A, Harvill ET. Evidence for horizontal gene transfer of two antigenically distinct O antigens in Bordetella bronchiseptica. Infect Immun 2009;77:3249–57. https://doi.org/10.1128/IAI.01448-08.

[23] Bart MJ, Harris SR, Advani A, Arakawa Y, Bottero D, Bouchez V, et al. Global Population Structure and Evolution of Bordetella pertussis and Their Relationship with Vaccination. MBio 2014;5:e01074–14. https://doi.org/10.1128/mBio.01074-14.

[24] Xu Z, Wang Z, Luan Y, Li Y, Liu X, Peng X, et al. Genomic epidemiology of erythromycin-resistant Bordetella pertussis in China. Emerg Microbes Infect 2019;8:461–70. https://doi.org/10.1080/22221751.2019.1587315.

[25] Feng Y, Chiu C-H, Heininger U, Hozbor DF, Tan TQ, von König C-HW. Emerging macrolide resistance in Bordetella pertussis in mainland China: Findings and warning from the global pertussis initiative. Lancet Reg Health West Pac 2021;8:100098. https://doi.org/10.1016/j.lanwpc.2021.100098.

[26] Zomer A, Otsuka N, Hiramatsu Y, Kamachi K, Nishimura N, Ozaki T, et al. Bordetella pertussis population dynamics and phylogeny in Japan after adoption of acellular pertussis vaccines. Microb Genomics 2018;4. https://doi.org/10.1099/mgen.0.000180.

[27] Ma L, Caulfield A, Dewan KK, Harvill ET. Pertactin-Deficient Bordetella pertussis, Vaccine-Driven Evolution, and Reemergence of Pertussis. Emerg Infect Dis 2021;27:1561–6. https://doi.org/10.3201/eid2706.203850.

[28] Jolley KA, Bray JE, Maiden MCJ. Open-access bacterial population genomics: BIGSdb software, the PubMLST.org website and their applications. Wellcome Open Res 2018;3:124. https://doi.org/10.12688/wellcomeopenres.14826.1.

[29] Bouchez V, Guglielmini J, Dazas M, Landier A, Toubiana J, Guillot S, et al. Genomic Sequencing of Bordetella pertussis for Epidemiology and Global Surveillance of Whooping Cough. Emerg Infect Dis 2018;24:988–94. https://doi.org/10.3201/eid2406.171464.

[30] Criscuolo A, Brisse S. AlienTrimmer: a tool to quickly and accurately trim off multiple short contaminant sequences from high-throughput sequencing reads. Genomics 2013;102:500–6. https://doi.org/10.1016/j.ygeno.2013.07.011.

[31] Liu Y, Schröder J, Schmidt B. Musket: a multistage k-mer spectrum-based error corrector for Illumina sequence data. Bioinforma Oxf Engl 2013;29:308–15. https://doi.org/10.1093/bioinformatics/bts690.

[32] Magoč T, Salzberg SL. FLASH: fast length adjustment of short reads to improve genome assemblies. Bioinforma Oxf Engl 2011;27:2957–63. https://doi.org/10.1093/bioinformatics/btr507.

[33] Crusoe MR, Alameldin HF, Awad S, Boucher E, Caldwell A, Cartwright R, et al. The khmer software package: enabling efficient nucleotide sequence analysis. F1000Research 2015;4:900. https://doi.org/10.12688/f1000research.6924.1.

[34] Bankevich A, Nurk S, Antipov D, Gurevich AA, Dvorkin M, Kulikov AS, et al. SPAdes: a new genome assembly algorithm and its applications to single-cell sequencing. J Comput Biol J Comput Mol Cell Biol 2012;19:455–77. https://doi.org/10.1089/cmb.2012.0021.

[35] Maiden MCJ, Jansen van Rensburg MJ, Bray JE, Earle SG, Ford SA, Jolley KA, et al. MLST revisited: the gene-by-gene approach to bacterial genomics. Nat Rev Microbiol 2013;11:728–36. https://doi.org/10.1038/nrmicro3093.

[36] Bratcher HB, Corton C, Jolley KA, Parkhill J, Maiden MCJ. A gene-by-gene population genomics platform: de novo assembly, annotation and genealogical analysis of 108 representative Neisseria meningitidis genomes. BMC Genomics 2014;15:1138. https://doi.org/10.1186/1471-2164-15-1138.

[37] Ka J, Mc M. BIGSdb: Scalable analysis of bacterial genome variation at the population level. BMC Bioinformatics 2010;11:595–595. https://doi.org/10.1186/1471-2105-11-595.

[38] Parkhill J, Sebaihia M, Preston A, Murphy LD, Thomson N, Harris DE, et al. Comparative analysis of the genome sequences of Bordetella pertussis, Bordetella parapertussis and Bordetella bronchiseptica. Nat Genet 2003;35:32–40. https://doi.org/10.1038/ng1227.

[39] Zhang S, Xu Y, Zhou Z, Wang S, Yang R, Wang J, et al. Complete genome sequence of Bordetella pertussis CS, a Chinese pertussis vaccine strain. J Bacteriol 2011;193:4017–8. https://doi.org/10.1128/JB.05184-11.

[40] Sebaihia M, Preston A, Maskell DJ, Kuzmiak H, Connell TD, King ND, et al. Comparison of the Genome Sequence of the Poultry Pathogen Bordetella avium with Those of B. bronchiseptica, B. pertussis, and B. parapertussis Reveals Extensive Diversity in Surface Structures Associated with Host Interaction. J Bacteriol 2006;188:6002–15. https://doi.org/10.1128/JB.01927-05.

[41] Guillot S, Descours G, Gillet Y, Etienne J, Floret D, Guiso N. Macrolide-Resistant Bordetella pertussis Infection in Newborn Girl, France. Emerg Infect Dis 2012;18:966–8. https://doi.org/10.3201/eid1806.120091.

[42] Bartkus JM, Juni BA, Ehresmann K, Miller CA, Sanden GN, Cassiday PK, et al. Identification of a mutation associated with erythromycin resistance in Bordetella pertussis: implications for surveillance of antimicrobial resistance. J Clin Microbiol 2003;41:1167–72. https://doi.org/10.1128/JCM.41.3.1167-1172.2003.

[43] van Loo IHM, Heuvelman KJ, King AJ, Mooi FR. Multilocus sequence typing of Bordetella pertussis based on surface protein genes. J Clin Microbiol 2002;40:1994–2001. https://doi.org/10.1128/JCM.40.6.1994-2001.2002.

[44] Croucher NJ, Page AJ, Connor TR, Delaney AJ, Keane JA, Bentley SD, et al. Rapid phylogenetic analysis of large samples of recombinant bacterial whole genome sequences using Gubbins. Nucleic Acids Res 2015;43:e15. https://doi.org/10.1093/nar/gku1196.

[45] Nguyen L-T, Schmidt HA, von Haeseler A, Minh BQ. IQ-TREE: A Fast and Effective Stochastic Algorithm for Estimating Maximum-Likelihood Phylogenies. Mol Biol Evol 2015;32:268–74. https://doi.org/10.1093/molbev/msu300.

[46] Letunic I, Bork P. Interactive Tree Of Life (iTOL) v4: recent updates and new developments. Nucleic Acids Res 2019;47:W256–9. https://doi.org/10.1093/nar/gkz239.

[47] Rambaut A. rambaut/figtree. 2021.

[48] Tazato N, Handa Y, Nishijima M, Kigawa R, Sano C, Sugiyama J 2015. Novel environmental species isolated from the plaster wall surface of mural paintings in the Takamatsuzuka tumulus: Bordetella muralis sp. nov., Bordetella tumulicola sp. nov. and Bordetella tumbae sp. nov. Int J Syst Evol Microbiol n.d.;65:4830–8. https://doi.org/10.1099/ijsem.0.000655.

[49] Yoon S-H, Ha S-M, Lim J, Kwon S, Chun J. A large-scale evaluation of algorithms to calculate average nucleotide identity. Antonie Van Leeuwenhoek 2017;110:1281–6. https://doi.org/10.1007/s10482-017-0844-4.

[50] Badhai J, Das SK. Genomic plasticity and antibody response of Bordetella bronchiseptica strain HT200, a natural variant from a thermal spring. FEMS Microbiol Lett 2021;368. https://doi.org/10.1093/femsle/fnab035.

[51] Kamanova J. Bordetella Type III Secretion Injectosome and Effector Proteins. Front Cell Infect Microbiol 2020;10:466. https://doi.org/10.3389/fcimb.2020.00466.

[52] Bayram J, Malcova I, Sinkovec L, Holubova J, Streparola G, Jurnecka D, et al. Cytotoxicity of the effector protein BteA was attenuated in Bordetella pertussis by insertion of an alanine residue. PLoS Pathog 2020;16:e1008512. https://doi.org/10.1371/journal.ppat.1008512.

[53] Buboltz AM, Nicholson TL, Parette MR, Hester SE, Parkhill J, Harvill ET. Replacement of adenylate cyclase toxin in a lineage of Bordetella bronchiseptica. J Bacteriol 2008;190:5502–11. https://doi.org/10.1128/JB.00226-08.

[54] Ahuja U, Liu M, Tomida S, Park J, Souda P, Whitelegge J, et al. Phenotypic and genomic analysis of hypervirulent human-associated Bordetella bronchiseptica. BMC Microbiol 2012;12:167. https://doi.org/10.1186/1471-2180-12-167.

[55] Weigand MR, Peng Y, Pouseele H, Kania D, Bowden KE, Williams MM, et al. Genomic Surveillance and Improved Molecular Typing of Bordetella pertussis Using wgMLST. J Clin Microbiol n.d.;59:e02726–20. https://doi.org/10.1128/JCM.02726-20.

[56] Porter JF, Connor K, Donachie W. Isolation and characterization of Bordetella parapertussis-like bacteria from ovine lungs. Microbiol Read Engl 1994;140 (Pt 2):255–61. https://doi.org/10.1099/13500872-140-2-255.

[57] Brinig MM, Register KB, Ackermann MR, Relman DA. Genomic features of Bordetella parapertussis clades with distinct host species specificity. Genome Biol 2006;7:R81. https://doi.org/10.1186/gb-2006-7-9-r81.

[58] Lund SJ, Rowe HA, Parton R, Donachie W. Adherence of ovine and human Bordetella parapertussis to continuous cell lines and ovine tracheal organ culture. FEMS Microbiol Lett 2001;194:197–200. https://doi.org/10.1111/j.1574-6968.2001.tb09469.x.

[59] Sealey KL, Belcher T, Preston A. Bordetella pertussis epidemiology and evolution in the light of pertussis resurgence. Infect Genet Evol J Mol Epidemiol Evol Genet Infect Dis 2016;40:136–43. https://doi.org/10.1016/j.meegid.2016.02.032.

[60] Barkoff A-M, Mertsola J, Pierard D, Dalby T, Hoegh SV, Guillot S, et al. Pertactin-deficient Bordetella pertussis isolates: evidence of increased circulation in Europe, 1998 to 2015. Euro Surveill Bull Eur Sur Mal Transm Eur Commun Dis Bull 2019;24. https://doi.org/10.2807/1560-7917.ES.2019.24.7.1700832.

[61] Bouchez V, Guillot S, Landier A, Armatys N, Matczak S, Toubiana J, et al. Evolution of Bordetella pertussis over a 23-year period in France, 1996 to 2018. Euro Surveill Bull Eur Sur Mal Transm Eur Commun Dis Bull 2021;26. https://doi.org/10.2807/1560-7917.ES.2021.26.37.2001213.

[62] Brito AF, Semenova E, Dudas G, Hassler GW, Kalinich CC, Kraemer MUG, et al. Global disparities in SARS-CoV-2 genomic surveillance. MedRxiv Prepr Serv Health Sci 2021. https://doi.org/10.1101/2021.08.21.21262393.

