## Supplementary File for "Genomic library of *Bordetella*"

**Contents :**

Supplementary figures	page 2

Supplementary tables	page 4

Supplementary text	page 5

Supplementary references	page 9

**Supplementary Figures**

**Figure S1: ANI heatmap of the *Bordetella* genus**

Heat map showing ANI values among Bordetella species, genogroups [1] and putative novel species. ANI values are colored from 75 (red) to 100 (blue). The heatmap was computed based on the ANI values matrix and ordered by internal classical hierarchical clustering of those values.

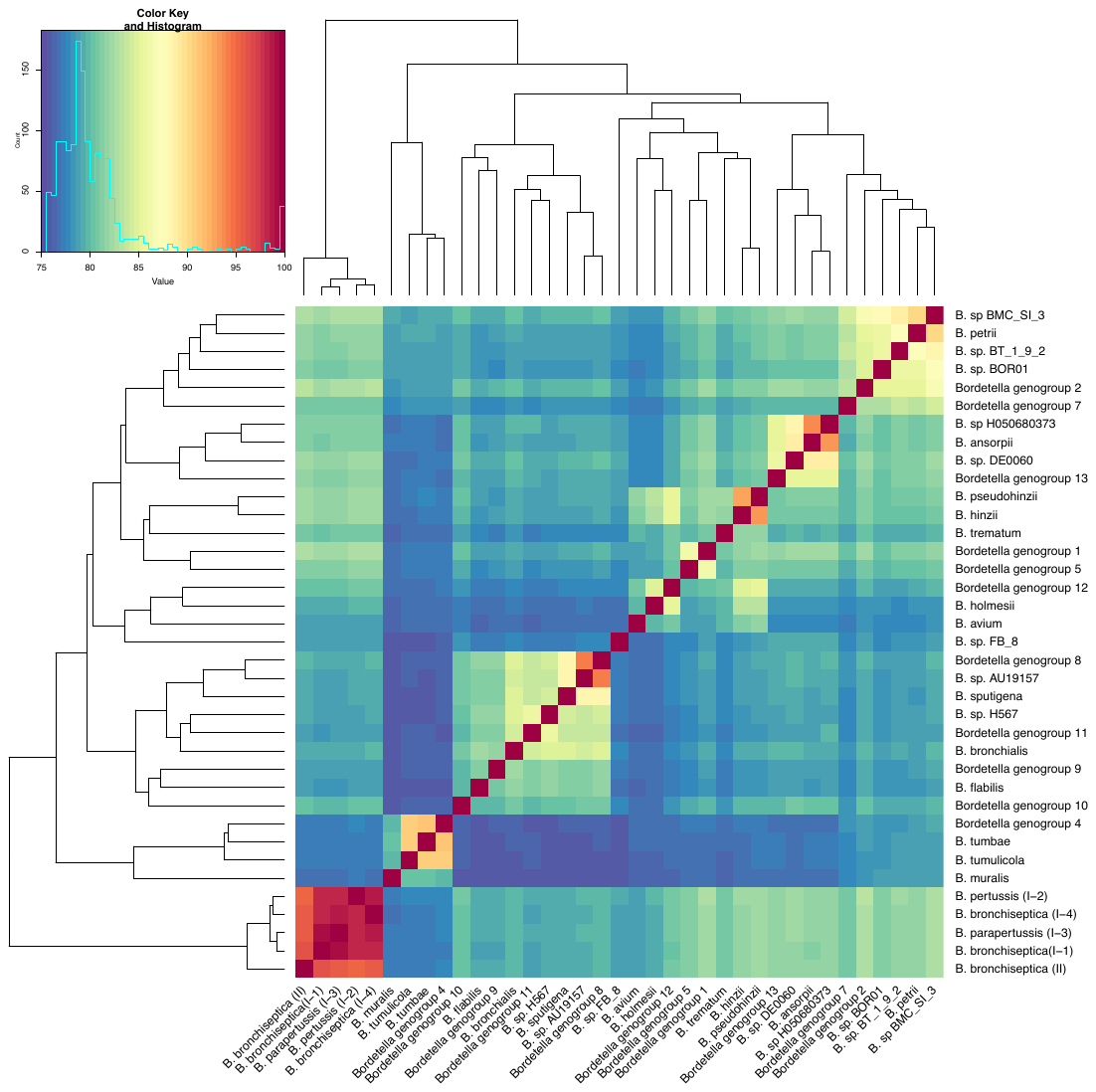

**Figure S2: Phylogeny of *B. bronchiseptica*** **with boostraps values of 100**

As for Figure 2, the analysis was performed with 186 *B. bronchiseptica* genomes, and representatives of the phylogenetic diversity of *B. pertussis* (9 genomes) and *B. parapertussis* (9 genomes). The recombination-purged concatenated multiple sequence alignment of 1,415 core gene loci (*cgMLST_genus* scheme) was used. The tree is rooted on lineage II, which is the most divergent clade. Isolates names are colored according to their lineage or sublineages (I-1, green; I-2, blue; I-3, red; I-4 brown; II, purple). Boostrap values equal to 100 are indicated on the branches by a blue circle.

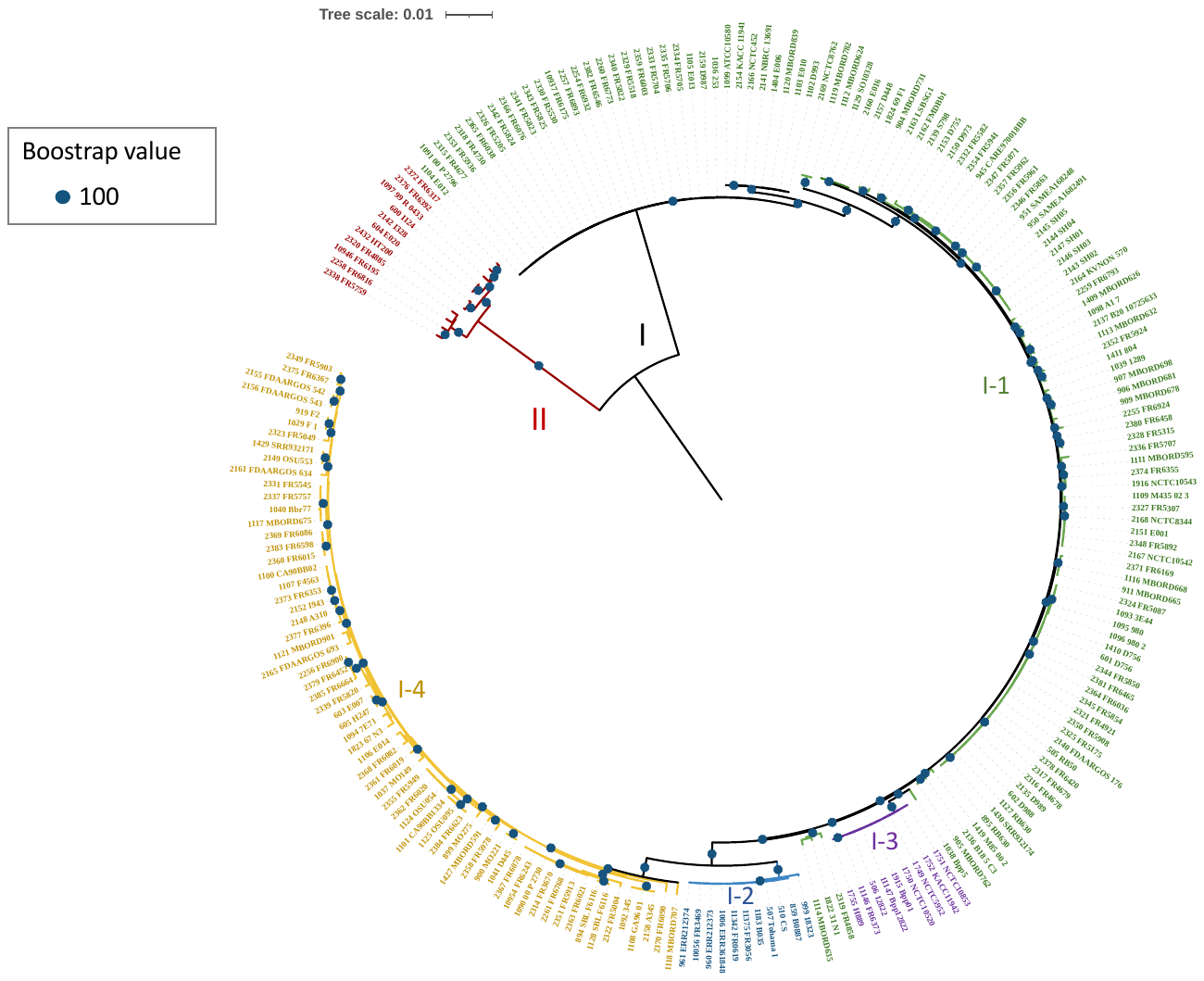

**Figure S3: Comparison of the allele call rates of two cgMLST schemes on classical *Bordetella* isolates*.***

The boxplots represent for each lineage or sublineage of the *B. bronchiseptica* genomic species (BbGS), the distribution of the rate of uncalled loci for the two cgMLST schemes: *cgMLST-pertussis* and *cgMLST-genus.* Boxplots are colored according to lineage or sublineages (I-1, blue; I-2, green; I-3, red; I-4, brown; II, purple).

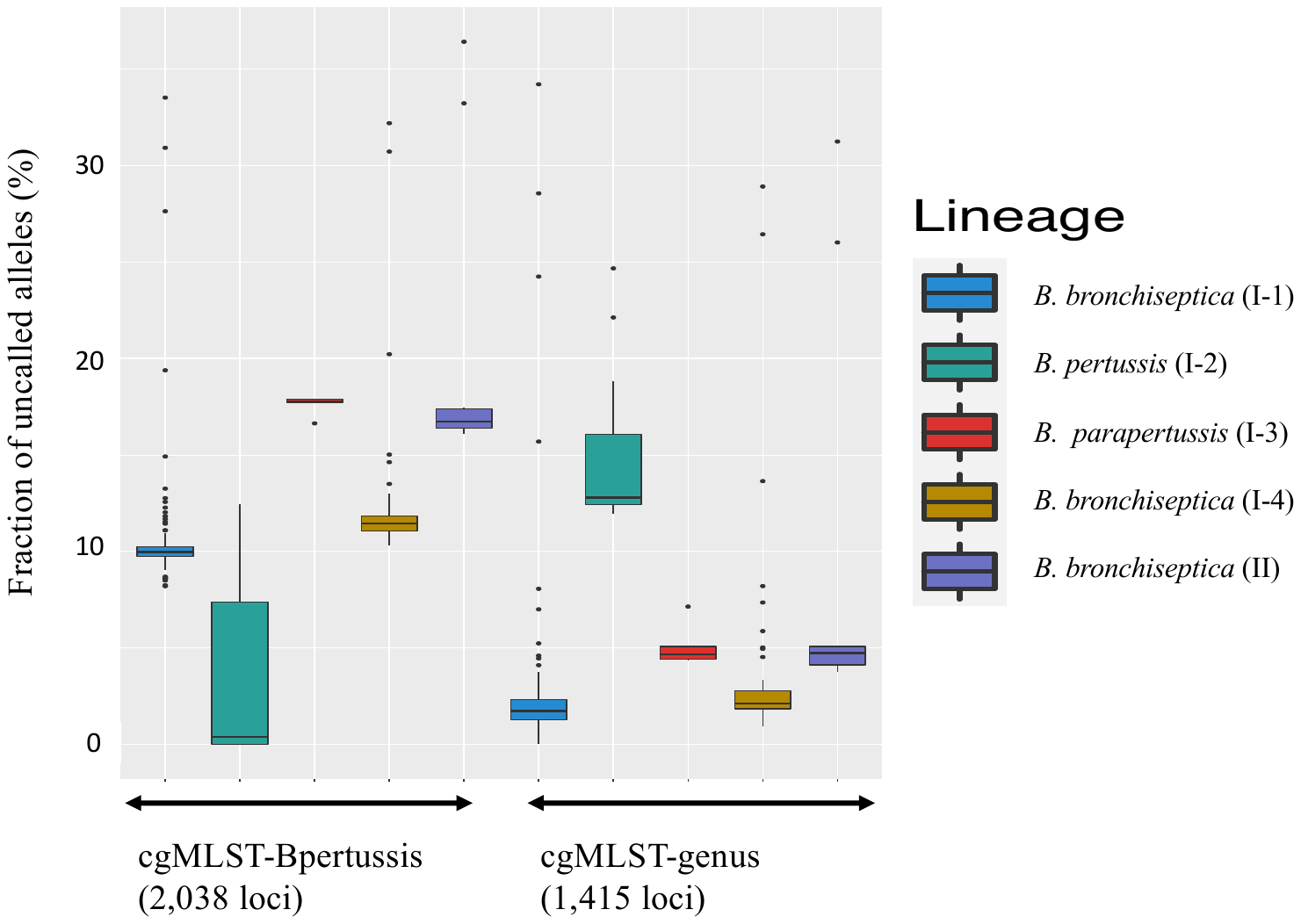

**Figure S4: Rooted phylogeny (built by JolyTree v2.1) of the *Bordetella* genus**

The phylogenetic tree was rooted using *Ralstonia* species as an outgroup. Bootstrap values are indicated using light blue circles (see key).

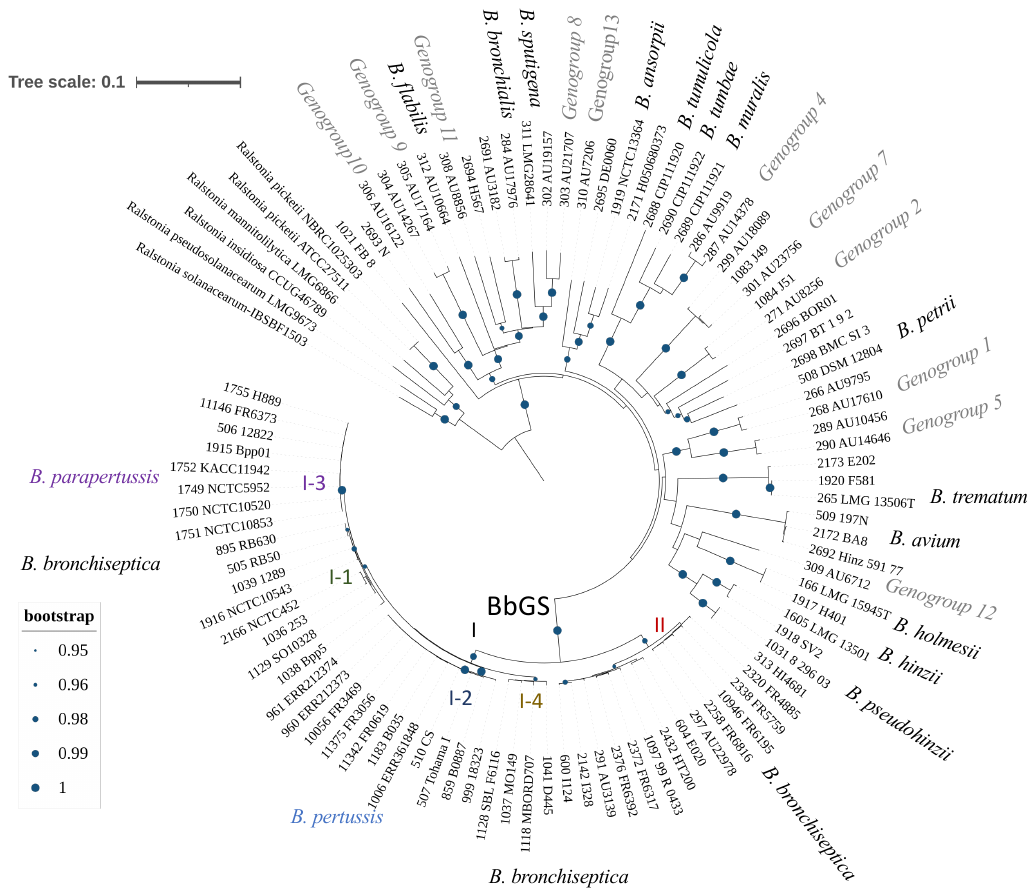

**Figure S5: Virulence allele profiles within the *Bordetella bronchiseptica* genomic species.** Same tree as in main Figure 2, with branch lengths. The outer circles represent allele variation at loci of the different virulence schemes, labeled by their BIGSdb scheme id: *Bp* Vaccine antigens (id 8), Phase biology genes (id 9), Other Toxins (10), T3SS (id 11), Autotransporters (id 12). Each color is specific to an allele number; across loci, similar color panels were used

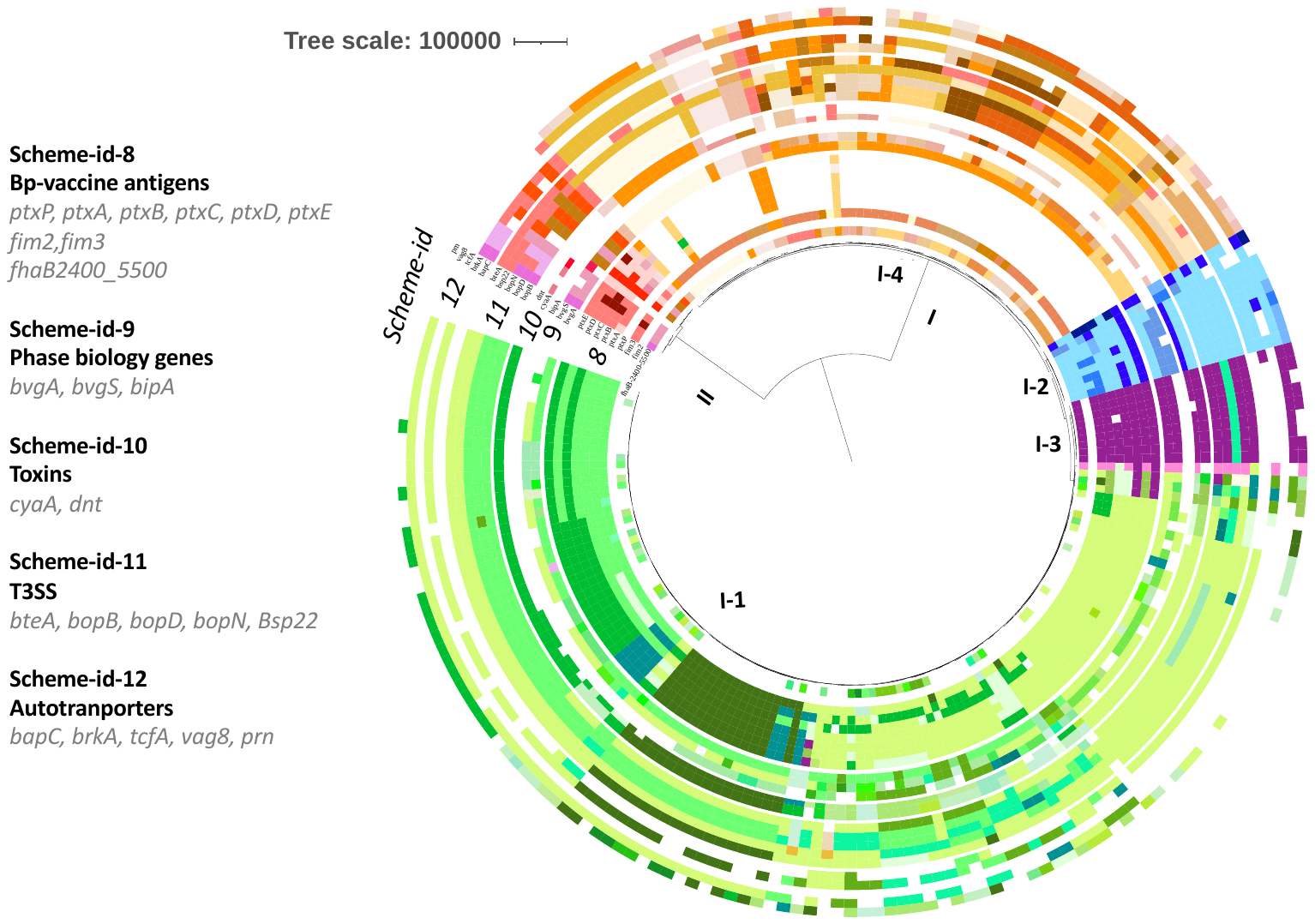
for each lineage or sublineage to illustrate lineage specificity of allele variation.

**Supplementary Tables**

**Supplementary Table 1: Correspondence between former alleles from Oxford BIGSdb and the current alleles in the merged database.**

**Supplementary Table 2: ANI values across the *Bordetella* genus.**

**Supplementary Table 3: Main alleles of the virulence related schemes observed in the BbGS lineages**

**Supplementary Table 4: *Bordetella* genomes list and accession numbers**

**Supplementary text: Variation of vaccine antigens and virulence-associated genes**

The main alleles of the loci of these schemes are summarized in **Supplementary Table 3**. We here describe the presence or absence of these genes, and the distribution of their alleles, within *B. bronchiseptica (Bbs)* and within *B. pertussis (Bp)*.

1. **Variation within *B. bronchiseptica***

Figure S5 provides a visual illustration of allele variation of the different genotyping schemes presented below.

***Bp-vaccine antigens* *scheme (fim2*, *fim3*, *ptxP*, *ptxA*, *ptxB*, *ptxC*, *ptxD*, *ptxE*, *fhaB-2400_5550)***

Even though pertussis toxin (PT) is known to be only produced by *Bp* isolates, the loci for PT promoter and subunits (*ptxP*, *ptxA*, *ptxB*, *ptxC*, *ptxD*, and *ptxE)* were present across most of the BbGS members. When present, alleles of these loci were specific for each lineage, *i.e.,* uniquely observed within a single lineage.

A notable exception was observed for 58 of 63 isolates from sublineage I-4, in which no alleles of the 6 PT-related loci were detected. This is in agreement with previous results underlying *ptxABCDE* absence in Bbr77 and MO149 isolates from sublineage I-4 [2]. Furthermore, we noticed that isolates of *Bbs* lineage II did not have alleles for the *ptxP* locus. This was true with the initial scan parameters (i.e. 90% identity, 90% alignment with the type allele) as well as when releasing these scan parameters to 50%. Regarding fimbriae, alleles were only detected in lineage I-2 (*Bp*) for locus *fim2* using our standard parameters. However, when releasing the scan parameters, we obtained partial fim2 matches for 78 isolates of lineages I-1, I-3, I-4 and 2, with identity percentage ranging from 78% to 97%.

For locus *fhaB* (2400-5550 region), no alleles were captured for 71 isolates, 67 of them belonging to sublineage I-1. Alleles were missing in 16/67 genomes but a partial match (from 1591 bp to 3099 bp over 3151 bp) was found when releasing scan parameters for the remaining isolates (51/67), with a high identity score (from 98 to 100%).

***Other toxins scheme***

For the two loci *cyaA* and *dnt*, included in the “Other toxins” scheme, we observed either sequence polymorphism or variation in the presence of these genes.

Gene *cyaA* encodes adenylate cyclase, an important toxin produced by isolates of the BbGS [3,4]. No alleles could be assigned for 32 isolates of lineages I-1 and 6 isolates of lineage II, consistent with previous findings: for strain 253 (BIGSdb ID: 1036) of lineage I-1 [4,5], the *cyaA* gene is replaced by a *ptp* operon [4]. Interestingly, in 5/6 isolates from lineage II (as for strains HT200 [6], BIGSdb ID: 2432), no alleles were found even when releasing the scan parameters.

The dermonecrotic toxin (DNT) is involved in turbinate atrophy and bronchopneumonia in pigs [7]. All isolates collected from pigs in our dataset were found in sublineage I-1 and had alleles at the *dnt* locus (as for example in strain S798 used to investigate DNT expression in Okada *et al* [8] (BIGSdb ID: 2139 ). However, we observed the absence of *dnt* alleles for isolates of lineage I-4, and for the ovine *B. parapertussis*, consistent with previous reports for lineage I-4 strains Bbr77 and D445 (BIGSdb ID 1040 and 1041) [5]. In the same way, strain HT200 of *Bbs* lineage II had no allele for *dnt* even when releasing scan parameters [6].

***Autotransporters scheme***

Autotransporters play an important role in *Bordetella* virulence. Among all autotransporters identified so far [9], five major autotransporters were included in the genotyping scheme: *prn*, *bapC, brkA, tcfA and vag8*.

Pertactin (Prn) is an important adhesin in the BbGS. This protein contributes to *Bbs* shedding and transmission between hosts [10]. All circulating BbGS isolates produced Prn before the use of vaccines, with a high variability among lineages due to different numbers of short repeats [11,12]. Since the introduction of acellular vaccines, an increasing number of isolates of lineages I-2 (*Bp*) and I-3 (*Bpp*) are deficient for Prn production [10,13]. This is reflected in our scanning results (Figure S5).

BapC (Bordetella Autotransporter Protein C) and BrkA (Bordetella Resistance to killing protein A) play a role in protecting *B. pertussis* from serum killing by complement [14,15]. Alleles were missing for *bapC* in many isolates of lineages I-1 and in all lineage I-3 (*Bpp*) isolates. In the same way, *brkA* alleles were missing in few isolates from lineage I-1 and in 8 of 10 isolates of lineage I-3 (*Bpp*).

Gene *tcfA*, coding for the tracheal colonization factor A, was only tagged for 6 of 11 isolates of sublineage I-2 (*Bp)*, consistent with literature [2,16,17].

Vag8 is an additional autotransporter involved in complement evasion [18]. Alleles were captured for isolates of lineages I-4 and I-2 (*Bp*) but were missing in some isolates of lineage I-1 and in all isolates of lineage II.

***T3SS scheme***

The allele variation at loci of this scheme is defined in the main text.

***Phase biology genes scheme***

Phase variation due to mutations in either *bvgA* or *bvgS* is frequent in *Bbs*. We found that *bvgA* and *bvgS* alleles were highly specific for each lineage or sublineage.

*bipA* sequence variation was described within the BbGS [19]. Even though *bipA* genes of *Bp* Tohama I and *Bbs* RB50 differ in the number of 90-amino-acid repeats and in their C-terminal sequence [19], *bipA* allele 3 was observed for isolates from Bp (sublineage I-2, comprising Tohama) and other alleles for some of sublineage I-1 (comprising RB50).

**2. Variation within *B. pertussis***

As expected, the diversity of alleles of the different schemes was much lower for *B. pertussis* than within the BbGS.

***Bp-vaccine antigen* *scheme (fim2*, *fim3*, *ptxP*, *ptxA*, *ptxB*, *ptxC*, *ptxD*, *ptxE*, *fhaB-2400_5550)***

The allele variation at loci of this scheme is defined in the main text.

***Other toxins scheme***

*Bp* isolates were characterized by *cyaA* allele 4*,* except for one isolate (ID 1240_B096) with allele 43. Eight isolates had no alleles but presented a partial match of 3402 to 3424 bp vs 5121 as expected with 100% identity to *cyaA* gene.

Regarding the *dnt* locus, Bp isolates had allele 1, except for 3 isolates with allele 4 (IDs 532, 533 and 10408) and another with allele 34 (ID 2089). Isolate FR6006 (ID 10220) had no allele for *dnt*.

***Autotransporters scheme***

Two *prn* alleles were predominant (alleles 1 and 2) in our dataset; as we matched our allele nomenclature with denominations present in the literature, this is consistent with the well-known shift from prn1 to prn2 that arose in the WCV period and increased in frequency in the ACV period [20]. We also detected 3 isolates with different alleles (prn-3, 7 and 9). In addition, because of the high prevalence of PRN-deficient isolates, often due to an incomplete *prn* CDS, a large proportion of isolates had no allele for *prn*, and these were mostly *ptxP3*.

Autotransporters genes *bapC* and *vag8* were highly conserved, as all *Bp* isolates were characterized by allele 1 for *bapC* locus and allele 4 for *vag8*.

Most Bp isolates had allele 1 for *brkA*, except for one isolate with allele 64 (ID 12189, isolate B1816); 14 isolates had no called allele.

*tcfA* was more diverse: although most isolates had allele 2, six other alleles were found, in one isolate each (alleles 4, 5, 11, 12, 13 and 35) and nine isolates had no allele. *Bp* isolates with no *tcfA* because of the entire gene deletion have previously been reported [17].

***T3SS scheme***

T3SS loci were highly conserved, and as a consequence T3ST-3 was found for most isolates. The main allele for *bopB, bopD* and *bopN*, *bsp22* or *betA* loci was allele 1. Three isolates had allele 4 for *bopB* and one had allele 33 for *bteA*. In addition, 11 isolates had no allele for *bteA*. *Bp* isolates producing no *bteA* have already been evidenced [21], such as FR0145 (ID 695) due to the insertion of an IS*481* in its promoter region.

***Phase biology genes scheme***

Almost all *Bp* isolates had allele 1 for *bvgA* and allele 5 for *bvgS.* One isolate presented a different *bvgA* allele (IDs 10140=FR4930, allele 2). In the same way, 5 isolates had a different *bvgS* allele (alleles 4, 6, 10, 11 and 63).

*bipA* allele was 3 in all *Bp* isolates and other alleles were captured for some *Bbs* isolates of sublineage I-1 (comprising RB50) consistent with previous findings [22].

**Supplementary references**

1.	Spilker T, Leber AL, Marcon MJ, et al. A simplified sequence-based identification scheme for Bordetella reveals several putative novel species. J Clin Microbiol **2014**; 52:674–677.

2.	Park J, Zhang Y, Buboltz AM, et al. Comparative genomics of the classical Bordetella subspecies: the evolution and exchange of virulence-associated diversity amongst closely related pathogens. BMC Genomics **2012**; 13:545.

3.	Chenal-Francisque V, Caro V, Boursaux-Eude C, Guiso N. Genomic analysis of the adenylate cyclase-hemolysin C-terminal region of Bordetella pertussis, Bordetella parapertussis and Bordetella bronchiseptica. Res Microbiol **2009**; 160:330–336.

4.	Buboltz AM, Nicholson TL, Parette MR, Hester SE, Parkhill J, Harvill ET. Replacement of adenylate cyclase toxin in a lineage of Bordetella bronchiseptica. J Bacteriol **2008**; 190:5502–5511.

5.	Park J, Zhang Y, Buboltz AM, et al. Comparative genomics of the classical Bordetella subspecies: the evolution and exchange of virulence-associated diversity amongst closely related pathogens. BMC Genomics **2012**; 13:545.

6.	Badhai J, Das SK. Genomic plasticity and antibody response of Bordetella bronchiseptica strain HT200, a natural variant from a thermal spring. FEMS Microbiol Lett **2021**; 368.

7.	Brockmeier SL, Register KB, Magyar T, Lax AJ, Pullinger GD, Kunkle RA. Role of the dermonecrotic toxin of Bordetella bronchiseptica in the pathogenesis of respiratory disease in swine. Infect Immun **2002**; 70:481–490.

8.	Okada K, Abe H, Ike F, et al. Polymorphisms influencing expression of dermonecrotic toxin in Bordetella bronchiseptica. PLoS One **2015**; 10:e0116604.

9.	Parkhill J, Sebaihia M, Preston A, et al. Comparative analysis of the genome sequences of Bordetella pertussis, Bordetella parapertussis and Bordetella bronchiseptica. Nat Genet **2003**; 35:32–40.

10.	Ma L, Dewan KK, Taylor-Mulneix DL, et al. Pertactin contributes to shedding and transmission of Bordetella bronchiseptica. PLoS Pathog **2021**; 17:e1009735.

11.	Boursaux-Eude C, Guiso N. Polymorphism of repeated regions of pertactin in Bordetella pertussis, Bordetella parapertussis, and Bordetella bronchiseptica. Infect Immun **2000**; 68:4815–4817.

12.	Diavatopoulos DA, Hijnen M, Mooi FR. Adaptive evolution of the Bordetella autotransporter pertactin. J Evol Biol **2006**; 19:1931–1938.

13.	Bouchez V, Brun D, Dore G, Njamkepo E, Guiso N. Bordetella parapertussis isolates not expressing pertactin circulating in France. Clin Microbiol Infect **2011**; 17:675–682.

14.	Bokhari H, Said F, Syed MA, et al. Molecular typing of Bordetella parapertussis isolates circulating in Pakistan. FEMS Immunol Med Microbiol **2011**; 63:373–380.

15.	Barnes MG, Weiss AA. BrkA protein of Bordetella pertussis inhibits the classical pathway of complement after C1 deposition. Infect Immun **2001**; 69:3067–3072.

16.	Finn TM, Stevens LA. Tracheal colonization factor: a Bordetella pertussis secreted virulence determinant. Mol Microbiol **1995**; 16:625–634.

17.	van Gent M, Pierard D, Lauwers S, van der Heide HGJ, King AJ, Mooi FR. Characterization of Bordetella pertussis clinical isolates that do not express the tracheal colonization factor. FEMS Immunol Med Microbiol **2007**; 51:149–154.

18.	Marr N, Shah NR, Lee R, Kim EJ, Fernandez RC. Bordetella pertussis autotransporter Vag8 binds human C1 esterase inhibitor and confers serum resistance. PLoS One **2011**; 6:e20585.

19.	Fuchslocher B, Millar LL, Cotter PA. Comparison of bipA alleles within and across Bordetella species. Infect Immun **2003**; 71:3043–3052.

20.	Bart MJ, Harris SR, Advani A, et al. Global population structure and evolution of Bordetella pertussis and their relationship with vaccination. MBio **2014**; 5:e01074.

21.	Hegerle N, Rayat L, Dore G, Zidane N, Bedouelle H, Guiso N. In-vitro and in-vivo analysis of the production of the Bordetella type three secretion system effector A in Bordetella pertussis, Bordetella parapertussis and Bordetella bronchiseptica. Microbes Infect **2013**; 15:399–408.

22.	Fuchslocher B, Millar LL, Cotter PA. Comparison of bipA alleles within and across Bordetella species. Infect Immun **2003**; 71:3043–3052.
